# Escape Excel: a tool for preventing gene symbol and accession conversion errors

**DOI:** 10.1101/103820

**Authors:** Eric A. Welsh, Paul A. Stewart, Brent M. Kuenzi

## Abstract

**Background:** Microsoft Excel automatically converts certain gene symbols, database accessions, and other alphanumeric text and numbers into dates, scientific notation, and other numerical representations, which may lead to subsequent, irreversible corruption of the imported text. A recent survey of popular genomic literature estimates that one-fifth of all papers with supplementary data containing gene lists in Excel format suffer from this issue.

**Results:** Here, we present an open-source tool, Escape Excel, which prevents these erroneous conversions by generating an escaped text file that can be safely imported into Excel. Escape Excel is available in the Galaxy web environment and can be installed through the Galaxy ToolShed. Escape Excel is also available as a stand-alone, command line Perl script on GitHub (http://www.github.com/pstew/escape_excel). A Galaxy test server implementation is accessible at http://apostl.moffitt.org.

**Conclusions:** Escape Excel detects and escapes a wide variety of problematic text strings so that they are not erroneously converted into other representations upon importation into Excel. Examples of problematic strings include date-like strings, time-like strings, leading zeroes in front of numbers, and long numeric and alpha-numeric identifiers that should not be automatically converted into scientific notation. It is hoped that greater awareness of these potential data-corruption issues, together with diligent escaping of text files prior to importation into Excel, will help to reduce the amount of Excel-corrupted data in scientific analyses and publications.

## Article

### Background

Auto-conversion of gene symbols and database accessions by the spreadsheet software Excel (Microsoft, Redmond WA) is a persistent issue in biomedical research. The first published account dates back to 2004, where gene symbols for the Serpin family (SEPT1, SEPT2, etc.) and others were reported as automatically converted to date formats within the software (1-Sep, 2-Sep, etc.) [1]. Furthermore, Riken database accessions such as 2310009E13 were reported as converted to the floating-point format of 2.31E+13. A recent programmatic scan of literature revealed that nearly a thousand supplementary files are affected by gene symbol to date auto-conversions [2]. Additional auto-conversions that occur include subsets of numbers with forward slashes or hyphens (converted to date format), numbers with colons (converted to time format), numbers followed by an A or P (converted to time format), and numbers containing leading zeros (leading zeros are dropped) [3]. Examples problematic text strings are given in Table 1.

**Table 1.** Escaped vs. unescaped text import into Excel.

| Escaped | Unescaped |
| --- | --- |
| 00123456 | 123456 |
| 2310009E13 | 2.31E+19 |
| 1234567890123456789 | 1.23457E+18 |
| SEPT7 | 07-Sep |
| April 31 | Apr-31 |
| April 99 | Apr-99 |
| 3 P | 3:00 PM |
| =1+2 | 3 |

Example strings are given that are auto-converted by Excel when imported from un-escaped text files. The escaped text is protected and remains unconverted after import

In order to avoid unwanted auto-conversions, text files can be imported via the File->Open menu within the Excel program, which then guides the user through options that allow the user to decide which columns to treat as text and which to treat as other data types. However, this method is time consuming, becomes impractical for large numbers of columns, cannot handle columns of mixed data types, and does not allow for customization of any columns beyond the 256^th^ column. Additionally, there is no option to select column data types when a text file is opened via "double clicking", "drag and drop", or "Open With" from the Windows file explorer or desktop. Thus, an automated method of escaping a text file prior to import in Excel, so as to protect vulnerable fields from unwanted auto-conversion, is necessary.

## Implementation

According to Microsoft, auto-conversion in Excel can be prevented by manually placing a single quotation (') in front of an entry as an escape character to force the cell to be interpreted as text [3]. However, if this method is used to escape a field within a file prior to importing into Excel, then the resulting field will contain an extra leading single quote after importation that must be dealt with. We have found that escaping a field within double quotes inside of an equation, such as ="*text to escape*", is able to protect strings from auto-conversion on text file import.

Escape Excel scans a tab-delimited text file for entries that need to be escaped. These entries are identified using regular expressions. All fields are first cleaned by removing leading/trailing spaces, enclosing quotes, leading single quotes, and UTF-8 byte order marks. These can interfere with correct parsing of the escaped output field by Excel, as well as interfere with regular expression pattern matching. Entries that are identified as likely to undergo unwanted auto-conversion by Excel are then encapsulated in a text string within an equation (e.g., SEPT1 is escaped as ="SEPT1"). Starting the entry with an equals sign tells Excel to treat the field as an equation, and the quotes tell Excel that the equation consists solely of a string. Once Escape Excel is run on a file, the resulting output can be opened in Excel without auto-conversion issues.

## Results

We have released the above implementation as Escape Excel, a freely available tool written in Perl for escaping a text file so as to prevent auto-conversion of known problematic entries upon opening in Excel. Escape Excel is implemented in the Galaxy [4] web environment and as a command line based Perl script (see Availability of data and materials). Examples of both Excel-safe text strings and strings that are escaped by the software are provided in Table S1 (in Additional file 1).

## Discussion

Most text-related functions in Excel will function properly on the escaped text equations, however, some features, such as Text-to-Columns will require the escaped text to be copied and pasted as "Paste Values" to convert the equations back into regular text strings before they will work correctly. Although this additional step of converting the escaped text back into regular text is not frequently required, it would be desirable to escape the text in such a manner that the text-within-equations work-around is not required. One such possibility would be to output an XML file consisting of the escaped text, with problematic fields specified as type "text". Although this could be used to import the problematic fields directly as text without going through an equation intermediate, there are several disadvantages to XML file output. The size of the file may increase significantly due to additional XML markup and formatting, which may be problematic for files of several hundred megabytes. The file would also no longer be in the form of a spreadsheet, which may reduce human readability. Although Escape Excel is envisioned to be run just prior to importing the file into Excel, thus obviating the need for compatibility outside of Excel, XML output would decrease compatibility with other software tools, including traditional UNIX row- and column- based command line utilities such as cut, paste, head, and tail, as well as other software that accepts tab-delimited input. The addition of an option to output XML files is high on our list of features for future work.

## Conclusions

We have developed and implemented a method for preventing Excel from corrupting imported text files due to unwanted auto-conversions. After processing with the Escape Excel software, text files can be safely imported into Excel without auto-converting gene symbols into dates and other related auto-conversion silent data corruptions. This open source software is freely available, and will be of great benefit in safely using Excel to analyze scientific data while minimizing the risk of silent data corruption.

## Declarations

### Ethics approval and consent to participate

Not applicable.

## Consent for publication

Not applicable.

## Availability of data and materials

Source code for Escape Excel is available at http://www.github.com/pstew/escape_excel. A Galaxy test server implementation is accessible at http://apostl.moffitt.org. The original text file of example strings used for generating Supplementary Table 1 is included in the software distribution. The version of the Perl script current as of the date of manuscript submission is included in the online supplementary material.

## Authors’ contributions

EAW developed the algorithm and implemented the software in Perl. PAS and BMK implemented the web server in Galaxy. BMK consulted and investigated alternative solutions. EAW and PAS were major contributors in writing this manuscript. All authors read and approved the final manuscript.

## Acknowledgements

Not applicable.

## List of abbreviations

Universal Transformation Format-8 (UTF-8), Extensible Markup Language (XML), Cancer Center Support Grant (CCSG), National Cancer Institute (NCI), Eric A. Welsh (EAW), Paul A. Stewart (PAS), Brent M. Kuenzi (BMK)

## References

1. Zeeberg BR, Riss J, Kane DW, Bussey KJ, Uchio E, Linehan WM, Barrett JC, Weinstein JN. Mistaken identifiers: gene name errors can be introduced inadvertently when using Excel in bioinformatics. BMC Bioinformatics 2004, 5:80.

2. Ziemann M, Eren Y, El-Osta A. Gene name errors are widespread in the scientific literature. Genome Biol 2016, 17:177.

3. Microsoft: Text or number converted to unintended number format in Excel. Redmond, WA, Microsoft; 2015. [https://support.microsoft.com/en-us/kb/214233] Accessed 29/08/16

4. Afgan E, Baker D, van den Beek M, Blankenberg D, Bouvier D, Cech M, Chilton J, Clements D, Coraor N, Eberhard C,et al. The Galaxy platform for accessible, reproducible and collaborative biomedical analyses: 2016 update. Nucleic Acids Res 2016, 44:W3-W10.

